# Identification a Compact Promoter using a New Promoter Selection Strategy and Engineering Hybrid Pol II/III Enable Efficient Genome Editing in Human Retinal Ganglion Cells

**DOI:** 10.64898/2026.01.26.701896

**Authors:** Ping-Wu Zhang, Steven H Zhang, Yen-Yu Chang, Sean Li, Laura Fan, Weifeng Li, Yukan Duan, Jie Cheng, Casey J. Keuthan, Cynthia A. Berlinicke, Derek S Welsbie, Donald J Zack

## Abstract

Promoters and vectors are critical components of gene therapy, enabling the delivery and expression of therapeutic genes to correct both loss- and gain-of-function mutations. Adeno-associated virus (AAV) vectors are the leading platform for in vivo gene delivery; however, the widely used Streptococcus pyogenes Cas9 (SpCas9, 4.1 kb) approaches the AAV packaging limit of 4.7 kb. This constraint often necessitates dual-vector systems, which reduce therapeutic efficiency, or the use of smaller nucleases such as SaCas9 (3.2 kb) and AacCas12b (3.4 kb), which have lower PAM site frequencies. To enhance promoter selection for gene therapy applications, we developed a strategy to identify compact, cell-preferred RNA polymerase II (Pol II) promoters. Analysis of approximately 300 compact Pol II promoters revealed that exogenous expression levels in one cell type correlate more strongly with those in other cell types than with endogenous expression, underscoring the importance of exogenous expression efficiency in promoter selection. Using this approach, we identified a compact Pol II promoter #2 (Pro2, 133 bp) that drives robust transgene expression in human retinal ganglion cells (RGCs). To enable single-AAV delivery of SpCas9, we analyzed three commonly used Pol III promoters (H1, 7SK and U6) and determined their minimal functional lengths using a CRISPR/Cas9 reporter assay. We further engineered three compact hybrid Pol II/III promoters which combined pro2 with minimal H1, 7SK and U6 (276, 294, and 323 bp) capable of co-expressing SpCas9 and gRNA, enabling efficient genome editing in both transfected HEK293 cells (approaching 100%) and human RGCs (up to 55.9%) from human stem cell–derived retinal ganglion cells (RGCs). Together, these findings establish a framework for developing single-AAV CRISPR-based gene therapy strategies.

**Authors’ contributions:** PWZ and DJZ conceived the study, designed the experiments, performed data analysis and interpretation, and were the primary contributors to manuscript writing. STZ played a key role in data collection and correlation analysis. YYC, SL, LF, CJK, YD, CAB, JC, and DW contributed to the execution of essential experiments and subsequent data analysis. All authors have read and approved the final manuscript.

**Declaration of interests:** The authors declare no conflicts of interest.

## Introduction

Gene therapy requires highly efficient delivery systems, such as viral vectors, and cell-or tissue-specific expression of therapeutic genes. This specificity is largely governed by RNA polymerase II (Pol II) promoters with adequate transcriptional strength and cell-type–preferred activity, or by the delivery system itself. However, identifying such promoters remains challenging, as current strategies often rely on endogenous gene expression profiles, which do not necessarily predict promoter performance in transgenic contexts.

Promoters in eukaryotes are broadly classified based on the type of RNA polymerase they recruit RNA polymerase I (Pol I), II (Pol II), or III (Pol III). Pol II promoters drive the transcription of protein-coding genes by recruiting RNA polymerase II [1], while Pol III promoters direct the transcription of small non-coding RNAs, such as 5S rRNA, tRNAs, and type-3 RNAs like U6 snRNA [2]. Among Pol III promoters, U6, 7SK, and H1 are commonly used in eukaryotic systems to drive guide RNA (gRNA) expression—one of the two key components of the CRISPR-Cas9 genome editing system [3].

More than 80% of current gene therapy approaches utilize adeno-associated virus (AAV) vectors [4]. Optimal AAV-based delivery requires several key features: compactness, delivery efficiency, and specificity. Compact promoters are especially important because the packaging capacity of AAV vectors is limited to ∼4.7 kb [5]. Achieving cell-or tissue-specific expression is another critical challenge, as there are few well-established strategies for systematically identifying compact and functional promoters suitable for gene therapy applications.

In CRISPR/Cas9-mediated gene therapy, high-expression endogenous promoters are often used to drive Cas9 expression. However, this approach has limitations: (1) such promoters may be too long to include all necessary regulatory elements within the AAV packaging constraint; and (2) screening and validating these promoters can be time-consuming and inefficient. Therefore, identifying compact promoters with robust exogenous activity in target cells is essential for efficient and targeted genome editing.

A major limitation of current CRISPR/Cas9 gene therapy strategies is the size of the commonly used SpCas9 gene (∼4.1 kb), which leaves insufficient space in a single AAV vector for additional essential elements. As a result, most approaches rely on dual-AAV systems, which suffer from reduced efficiency due to the need for co-infection. A single AAV approach would enable the co-delivery of all necessary components—Cas9 and gRNA—into the same cell, enhancing editing efficiency and precision. To enable this, the ideal promoter system would combine both Pol II and Pol III functions in under 350 base pairs (bp), leaving room for the Cas9 gene and minimal regulatory sequences (∼250 bp for a terminator and gRNA scaffold).

To address these challenges, we aimed to identify compact Pol II promoters with cell-preferred activity, particularly for human retinal ganglion cells (RGCs). Bidirectional promoters, which have clearly defined boundaries and compact structure, were attractive candidates for this purpose. A total of 1,352 bidirectional promoters with transcription start sites (TSSs) separated by less than 1,000 bp are available [6].

Gene therapy involves delivering genetic material into target tissues or cells to achieve therapeutic gene expression. The eye is an attractive organ for gene therapy due to its immune-privileged status, accessibility for delivery, and the existence of over 350 inherited ocular diseases. Within the eye, the retina is the primary target for gene therapy [7].

Our initial strategy, which prioritized promoters based on endogenous gene expression, yielded suboptimal results. This led us to hypothesize that exogenous expressions driven by candidate promoters may not directly correlate with their endogenous expression. To test this, we analyzed previously published luciferase data from over 300 bidirectional promoters across four cell types, comparing exogenous activity with endogenous expression levels [6, 8]. This guided our selection of compact Pol II promoters.

In parallel, we pursued a Pol II–Pol III hybrid promoter strategy and used gene editing efficiency as a functional readout to optimize Pol III promoter size for gRNA expression, replacing traditional Northern blot analysis. In this setup, Cas9 was driven by the same Pol II promoter [9]. Using this approach, we identified three compact chimeric promoters capable of driving robust and cell-preferred exogenous expression for efficient genome editing in human RGCs.

## Methods

### Correlation analysis of exogenous and endogenous gene expression

Luciferase expression data from over 300 bidirectional promoters in four cell lines—HeLa (human cervical cancer), WI-38 (human primary fibroblasts), HT1080 (human fibrosarcoma), and mouse embryonic fibroblasts (MEFs; ATCC)—were obtained from Trinklein et al. [6]. Corresponding endogenous gene expression data (RNA-seq) for the same cell lines were collected from publicly available datasets [10–14]. To evaluate the relationship between endogenous and exogenous promoter activity, correlation analyses were conducted using R software. Correlations between either exogenous transgene expression (luciferase transfection expression driven by the same promoters) or endogenous gene expression (RNA-seq), or both exogenous and endogenous gene expression between the four cell lines were performed using R program. Pearson correlation coefficients (r) were used to assess the degree of linear association. The strength of correlation was categorized as follows:

Very high correlation: 0.9 – 1.0

High correlation: 0.7 – 0.9

Moderate correlation: 0.5 – 0.7

Low correlation: 0.3 – 0.5

Little or no correlation: < 0.3

### Pol II/III chimera promoter construction

Restriction enzymes BamHI and NheI were used to isolate a 2,906 bp fragment from the plasmid pAAV-MCS2 (Addgene #46954) and a 5,025 bp fragment encoding Cas9-P2A-eGFP from Addgene #110866. These two fragments were ligated to generate a 7,943 bp recombinant plasmid, termed pAgR-Cas9-eGFP, which served both as a Pol II promoter testing platform and as a backbone for chimera promoter cloning.

To construct the chimeric Pol II/III promoter, a BamHI–scaffold–gRNA–hybrid promoter–EcoRI fragment was inserted upstream of the Kozak–Cas9–NLS–P2A–eGFP cassette within the pAgR-Cas9-eGFP backbone. All DNA fragments were assembled using a Gibson Assembly Kit (New England Biolabs, USA), following the manufacturer’s protocol.

The final construct, pAgRNA-Cas9-eGFP, targets the genomic sequence of the DLK gene and enables site-specific genome editing in the presence of Cas9 expression. The guide RNA sequence used for targeting DLK was: 5′–TGTGGAGAGTACATCAGCTG–3′.

### Human embryonic stem Cell (hESC) maintenance and differentiation to RGCs

hESCs were maintained by clonal propagation in mTeSR™ Plus medium (STEMCELL Technologies, USA) on growth factor-reduced Matrigel-coated plates under standard culture conditions (5% CO_2_, 21% O_2_). Colonies were passaged by enzymatic dissociation using Accutase (Sigma-Aldrich, USA). To support the survival of single cells, mTeSR Plus medium was supplemented with 5 μM blebbistatin. Differentiation of hESCs into retinal ganglion cells (RGCs) was performed using our previously established protocol [15]. Briefly, hESCs were dissociated into single cells and plated onto Matrigel-coated plates at a density of 52,600 cells/cm^2^ in mTeSR Plus supplemented with 5 μM blebbistatin. This plating day was designated as day −1 (d−1). On day 0, the medium was completely replaced with N2B27 differentiation medium, consisting of a 1:1 mixture of DMEM/F12 and Neurobasal medium supplemented with 1% GlutaMAX™, 1% antibiotic-antimycotic, 1% N2 Supplement, and 2% B27 Supplement (all from Thermo Fisher Scientific, USA), marking the start of differentiation.

### Genomic DNA extraction and PCR amplification of gene editing region

Genomic DNA (gDNA) was extracted from cultured cell monolayers using the Quick-DNA Microprep Kit (Zymo Research, USA) following the manufacturer’s instructions. Briefly, HEK293A cells were detached using trypsin, pelleted by centrifugation in DMEM, and processed for gDNA isolation. Eluted DNA was quantified using a NanoDrop 100 spectrophotometer (Thermo Fisher Scientific, USA). PCR was performed in a 20-μL reaction volume consisting of 50ng genomic DNA, a final concentration of 250nm primers, and 10 μl 2X Phusion Flash PCR Master Mix (Thermo Fisher Scientific, USA). The following primer pair was used for PCR amplification of the DLK reporter gene: Forward primer: 5′-CTCTTCTAGGCGGTGCAGAC–3′; Reverse primer: 5′–GGACACACTTTGGGAAAGGA–3′. Amplification was carried out under the following conditions: 98°C for 2 min, followed by 38 cycles of 98°C for 10 s, 60°C for 10 s, and 72°C for 10 s, with a final extension at 72°C for 2 min. The PCR products were analyzed on 1.2% agarose gel using 1X TAE buffer containing 10 mg/mL ethidium bromide or Ultra GelRed (Vazyme, P. R. China). The amplified DNA fragments were subsequently used as templates for restriction enzyme digestion reactions.

### Digestion and DNA gel electrophoresis

PCR products were digested with PvuII-HF restriction enzyme to assess targeted modifications. Each digestion was performed in a 15 μl reaction containing: 1 μg template DNA, 1.5 μl 10X CutSmart Buffer (New England Biolabs, USA), and 15 U PvuII-HF (New England Biolabs, USA). Reactions were incubated at 37°C for either 2 hours or 24 hours to ensure complete digestion. Digestion products were resolved on 1.2% agarose gels in 1X TAE buffer, running at 100 V for 50 minutes. A 1 kb Plus DNA ladder (Thermo Fisher Scientific, USA) was used as a molecular size reference.

### Transfection of HEK293 cells and hESC-Derived RGCs

HEK293A cells were cultured in DMEM supplemented with 10% FBS and transfected at 30– 60% confluency using Lipofectamine™ 3000 Transfection Reagent (Thermo Fisher Scientific, USA) according to the manufacturer’s protocol. Plasmids containing candidate promoters were used for transfection. hESC-derived retinal ganglion cells (RGCs) and other neuron-related cells were transfected with plasmids harboring chimeric promoters using the Lipofectamine Stem Transfection Reagent Kit (Thermo Fisher Scientific, USA), following the supplier’s instructions.

### Gene editing analysis

If not specified all the gene editing analysis were carried out on a reporting gene DLK (MAP3K12) and HEK293 cell transfection (genomic DNA were collected 48h after transfection). A 509-base pair (bp) segment of the DLK reporter gene was PCR-amplified for gene editing analysis. This segment contains a PvuII restriction site (CAGCTG). In the absence of gene editing, digestion of the 509 bp PCR product with PvuII-HF yields two fragments of 358 bp and 151 bp. When gene editing occurs, the restriction site is disrupted, resulting in an undigested 509 bp fragment. Gene editing efficiency in HEK293A and hESC-derived retinal ganglion cells (RGCs) were quantified by calculating the ratio of the intact 509 bp fragment relative to the sum of all fragments (509 bp + 358 bp + 151 bp). Densitometry analysis was performed using Image Lab 6.0 software (Bio-Rad).

### FACS purification

HEK293A cells were dissociated into single-cell suspensions by treatment with 0.25% Trypsin-EDTA for 5 minutes, washed once with PBS, and resuspended in DMEM containing 10% FBS. hESC-derived RGCs and CRX-GFP cells were dissociated using AccuMAX for 45 minutes, washed once with PBS, and resuspended in their original culture medium. GFP-positive cells were enriched from 600 μl of cell suspension using a Sony SH800 cell sorter (Sony, Japan). During sorting, cell debris was excluded by forward scatter (FSC) and side scatter (SSC) gating. Doublet discrimination was performed by gating on cells with proportional FSC-A/FSC-H signals. Cells were excited with a 488 nm laser, and fluorescence was detected on the FL2 channel (525/50 nm). The GFP-positive population was gated based on a fluorescence threshold set using non-transfected control cells. Cells with fluorescence above this threshold were collected as transfected, while those below were collected as negative.

### Human stem cell-Derived retinal organoid differentiation and cell dissociation

H9 knock-in reporter embryonic stem cells (ESCs) expressing tdTomato under the control of the cone-rod homeobox (CRX) promoter (H9-CRX) were differentiated into 3D retinal organoids following a previously published protocol [16], with minor modifications. At day 80, retinal organoids (n = 12) were enzymatically dissociated into single cells using HBSS (Thermo Fisher Scientific, USA) containing 1 mg/mL papain suspension (Worthington Biochemical, USA). The papain was pre-activated according to the manufacturer’s instructions.

Organoids were incubated in the papain solution at 37°C in a water bath with gentle trituration every 10 minutes for approximately 1 hour to promote dissociation. The enzymatic reaction was then quenched by dilution in culture medium supplemented with 10% fetal bovine serum (FBS). Dissociated cells from all organoids were pooled and filtered through a 40 μm cell strainer (VWR, USA) before plating. Approximately 44,000 cells per well were seeded onto poly-D-lysine/laminin-coated 24-well plates or coverslips. Cells were cultured in retinal organoid medium composed of a 3:1 mixture of DMEM: F12, supplemented with:10% FBS,100 μM taurine (Sigma Aldrich, USA), 1% MEM non-essential amino acids (NEAA) solution, 1% antibiotic-antimycotic solution, 1% N2 supplement (all reagents from Thermo Fisher Scientific, USA except taurine). Cells were allowed to recover and attach for approximately 24 hours prior to transfection.

### Fluorescent immunocytochemistry of retinal organoid Cells

Transfected cells were fixed in 4% paraformaldehyde (PFA) in phosphate-buffered saline (PBS) for 20 minutes at room temperature (RT). Cells were then permeabilized and blocked with 0.25% Triton X-100 and 4% bovine serum albumin (BSA) in PBS for 30 minutes at RT.

Following blocking, cells were incubated overnight at 4°C with primary antibodies diluted in 3% BSA in PBS. The primary antibodies used included: Anti-RFP/tdTomato (ChromoTek 5F8, USA), Anti-GFP (Abcam, ab13970, USA), Anti-βIII Tubulin (BioLegend, 801201, USA). After primary incubation, cells were washed and incubated with species-specific secondary antibodies conjugated to fluorophores: Alexa Fluor 488 Donkey Anti-Chicken, Alexa Fluor 568 Donkey Anti-Rat, Alexa Fluor 405 Donkey Anti-Mouse (all from Invitrogen, USA). Cells were mounted using antifade mounting medium with or without DAPI (Vectashield, H-1200-10 or H-1000-10, USA). Fluorescent images were captured on a Zeiss LSM 710 confocal microscope system.

## Results

### Transgene Expression Driven by Compact Promoters Does Not Correlate with Endogenous Gene Expression

The spatiotemporal regulation of exogenous gene expression remains a major challenge in gene therapy. Traditionally, endogenous gene expression patterns have been used as a reference to guide promoter selection. However, whether exogenous transgene expression levels truly correlate with endogenous gene expression remains unclear.

To investigate this, we analyzed transgene expressions driven by over 300 compact bidirectional promoters (each <1 kb) across four different cell lines and compared these levels to the corresponding endogenous gene expression for each cell line. Following the nomenclature established by Trinklein et al. [6], the gene transcribed in the sense strand direction corresponds to the forward promoter, and the gene transcribed in the antisense strand corresponds to the reverse promoter.

Surprisingly, neither forward nor reverse transgene expression showed significant correlation with endogenous gene expression. All calculated Pearson correlation coefficients (r) were below 0.35, indicating a weak or negligible relationship between exogenous and endogenous expression levels (Figure 1, Table 1). According to the correlation ranges defined in the Methods, coefficients between 0.3 and 0.5 represent low correlation, while those below 0.3 indicate no correlation.

**Table 1.**
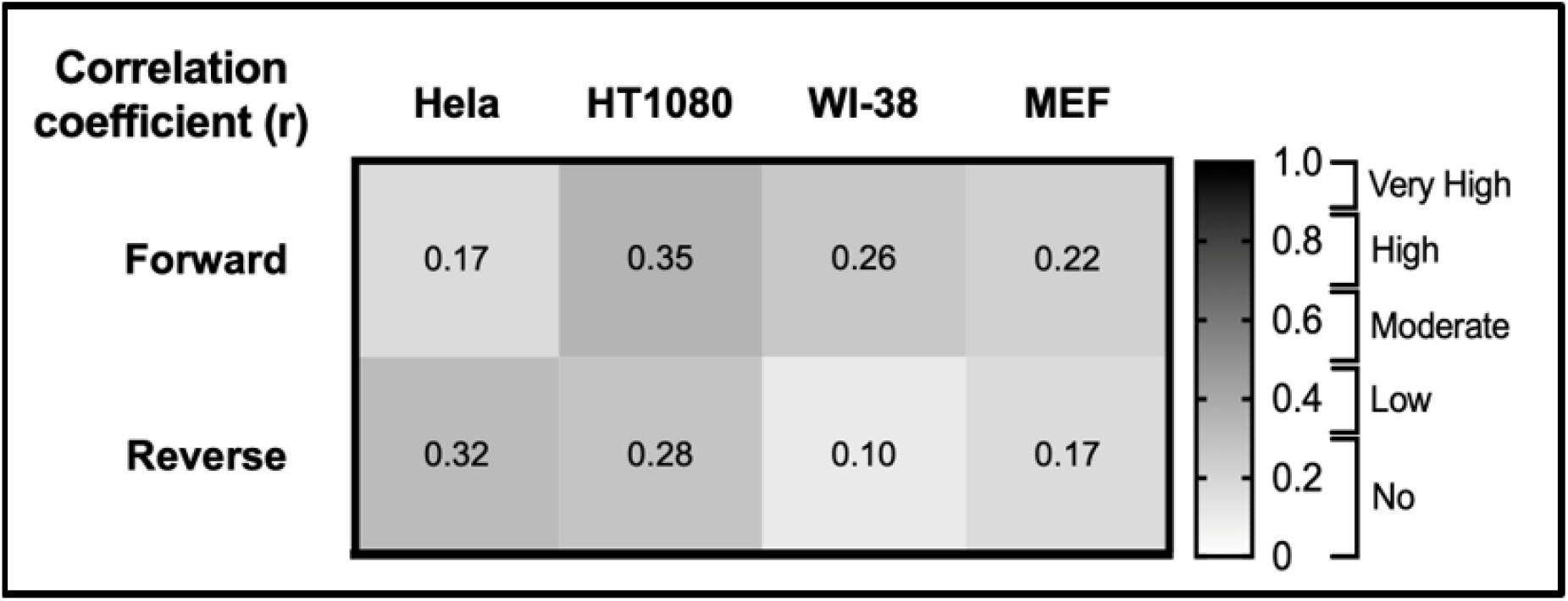
Correlation between exogenous and endogenous genes in four cell lines.

**Figure 1.**
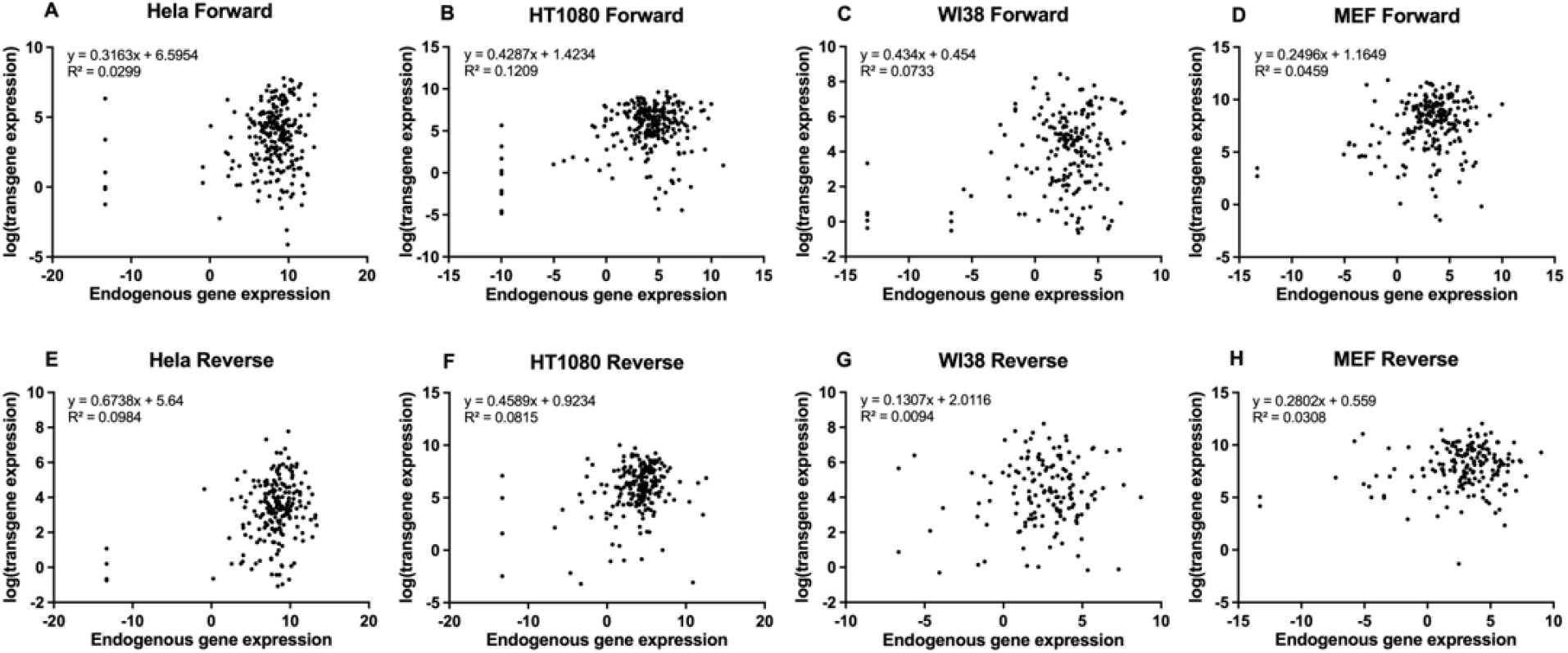
Low or no correlation between exogenous transgene and endogenous gene expression. Correlation analysis was performed to compare exogenous transgene expressions by luciferase reporter assays with endogenous gene expression data in four cell types (RNA-seq): Hela (a cervical cancer cell line), HT1080 (a fibrosarcoma line), WI-38 (primary human fibroblasts), and mouse embryonic fibroblasts (MEFs). R^2^ denotes the correlation of determination derived from the correlation coefficient(r), to indicate the strength of the linear relationship. A-D) Correlation analysis for forward promoters. E-H) Correlation analysis for forward promoters.

### Endogenous Gene Expression Correlates Across Cell Lines with Varying Strength

We next examined the extent to which endogenous gene expression levels correlate across the four cell lines. Forward promoter-driven expression showed low to moderate correlations among the human cell lines, while mouse embryonic fibroblasts (MEFs) exhibited low or negligible correlation with the others. In contrast, reverse promoter-driven gene expression displayed moderate to high correlations across all four cell lines. The strongest correlations were observed between WI-38 and HT1080 cells (R^2^ = 0.7663), and between HeLa and HT1080 cells (R^2^ = 0.4317). MEFs consistently showed weaker correlations with the other cell types, indicating more distinct expression profiles. These results highlight the cell-type specificity and variability of endogenous gene expression patterns (Figure 2, Table 2).

**Table 2.**
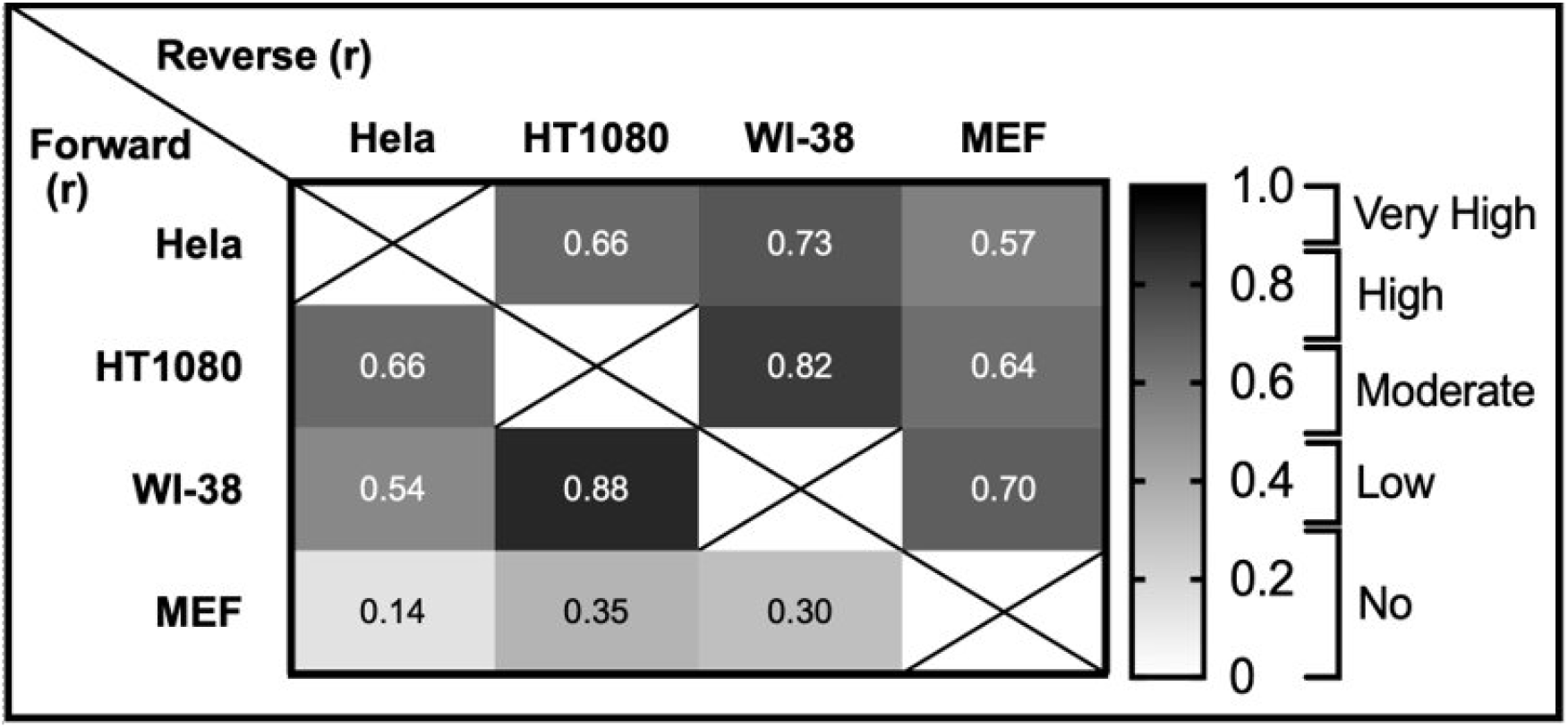
Correlation of endogenous genes among four cell lines.

**Figure 2.**
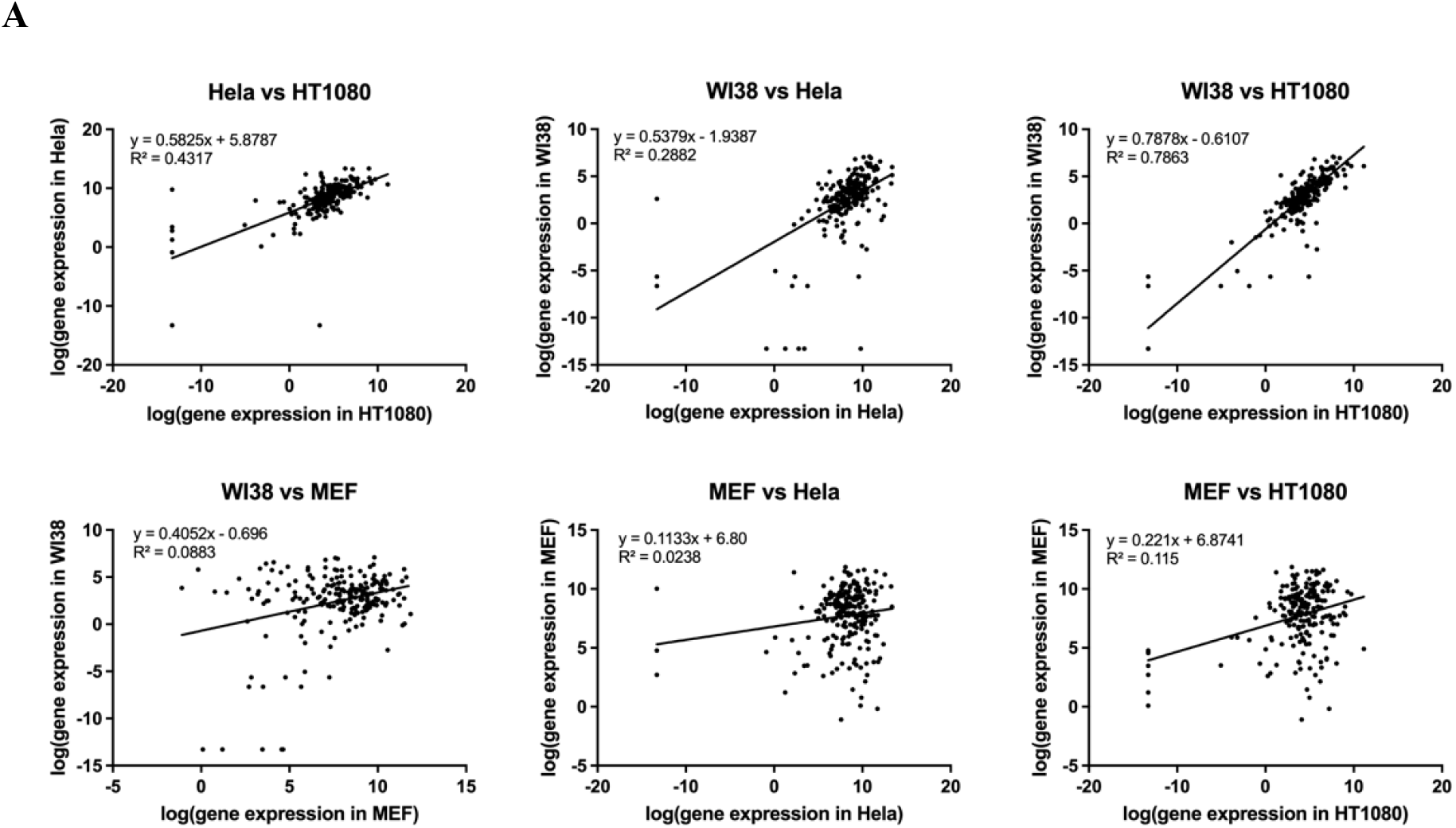
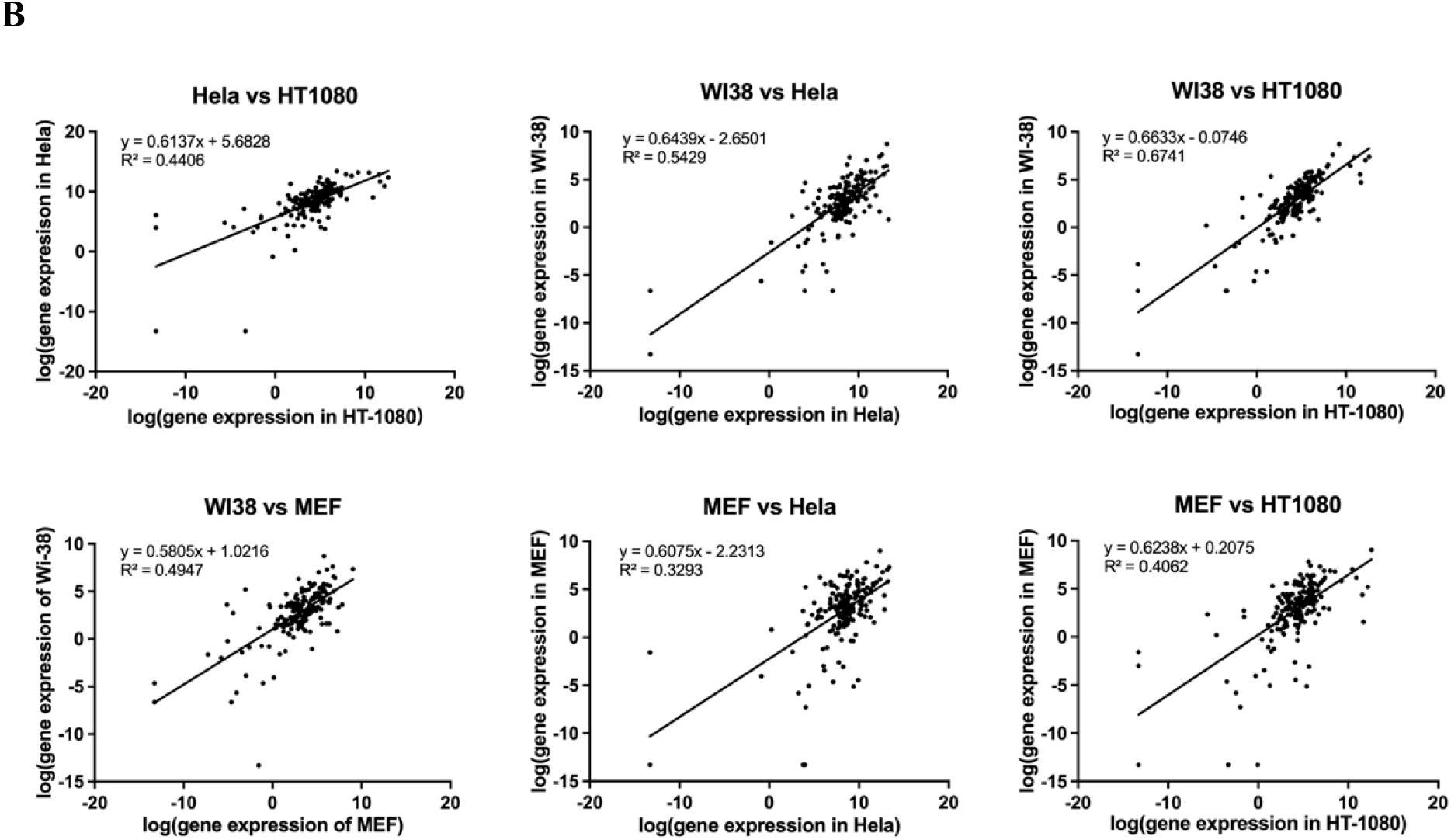
Correlations in endogenous gene expression can differ among cell lines. Scatter plots comparing log-transformed endogenous gene expression levels between pairs of cell lines: HeLa, HT1080, WI38, and MEF. Each dot represents the expression of a single gene measured in both cell lines of the pair. A) Comparison between forward promoters. B) Comparison between reverse promoters.

### Forward Promoter-Driven Exogenous Transgene Expression Shows Strong Cross-Cell Line Correlation

We next assessed the correlation of exogenous transgene expression driven by various promoters across different cell lines. Forward promoter-driven expression exhibited strong to very strong correlations among all cell lines, indicating consistent transgene regulation across cellular contexts. Reverse promoter-driven expression also showed generally high correlations, with the notable exception of WI-38 human primary fibroblasts, which displayed weak correlations with the other cell types. These findings suggest that while forward promoter activity is broadly conserved across cell types, reverse promoter-driven expression may be more sensitive to cell-type-specific regulatory environments (Fig. 3, Table 3).

**Table 3.**
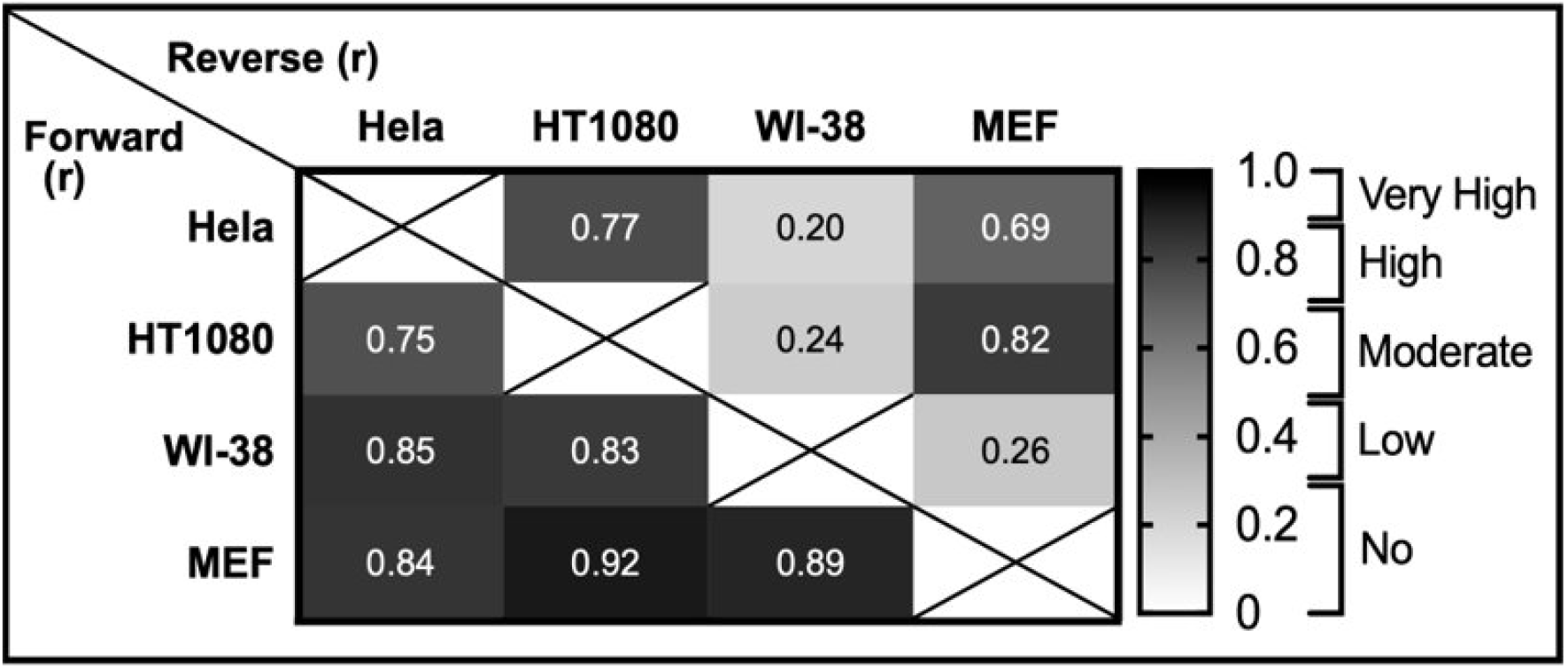
Correlation of exogenous genes among 4 cell lines.

**Figure 3.**
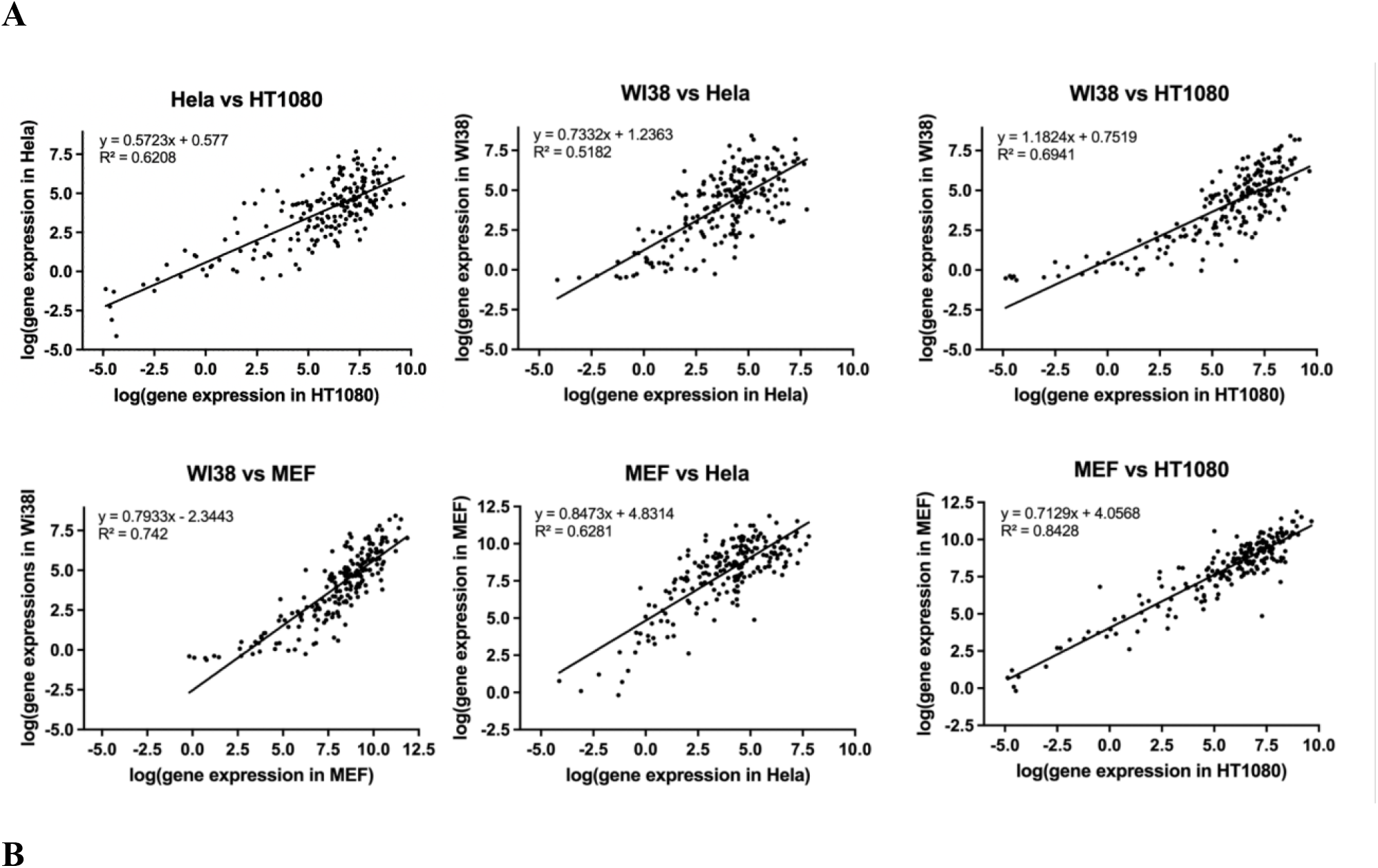
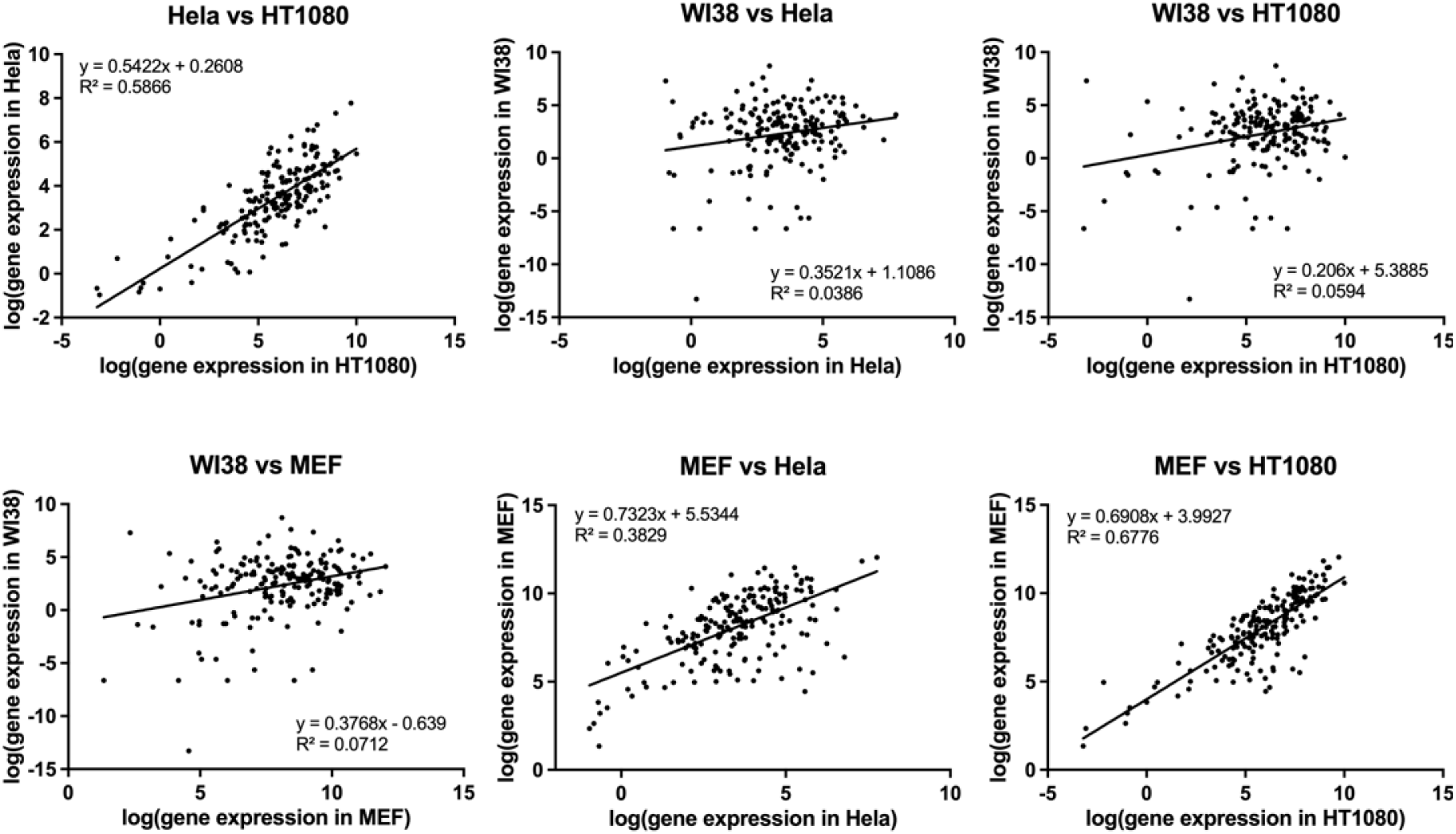
Forward exogenous genes showed high or very high correlation among 4 cell lines. A) Correlation comparison for forward promoters. B) Correlation comparison for reverse promoters.

### Screening for Compact Pol II Promoters Based on Exogenous Gene Expression

The human H1 and 7SK promoters possess both Pol II and Pol III promoter activity [17,19]. However, our experiments showed that the H1 promoter drives only minimal GFP expression in hESC-derived retinal ganglion cells (RGCs) (Figure 4B, 4C), and a similarly low level of GFP expression was observed with the 7SK promoter. Given the limited activity of naturally occurring dual Pol II–Pol III promoters in human RGCs, we aimed to engineer a chimeric promoter with both functions. As a first step, we screened a panel of bidirectional promoters to identify compact and efficient Pol II promoters capable of driving exogenous gene expression in hESC-derived RGCs (Figure 4A).

**Figure 4.**
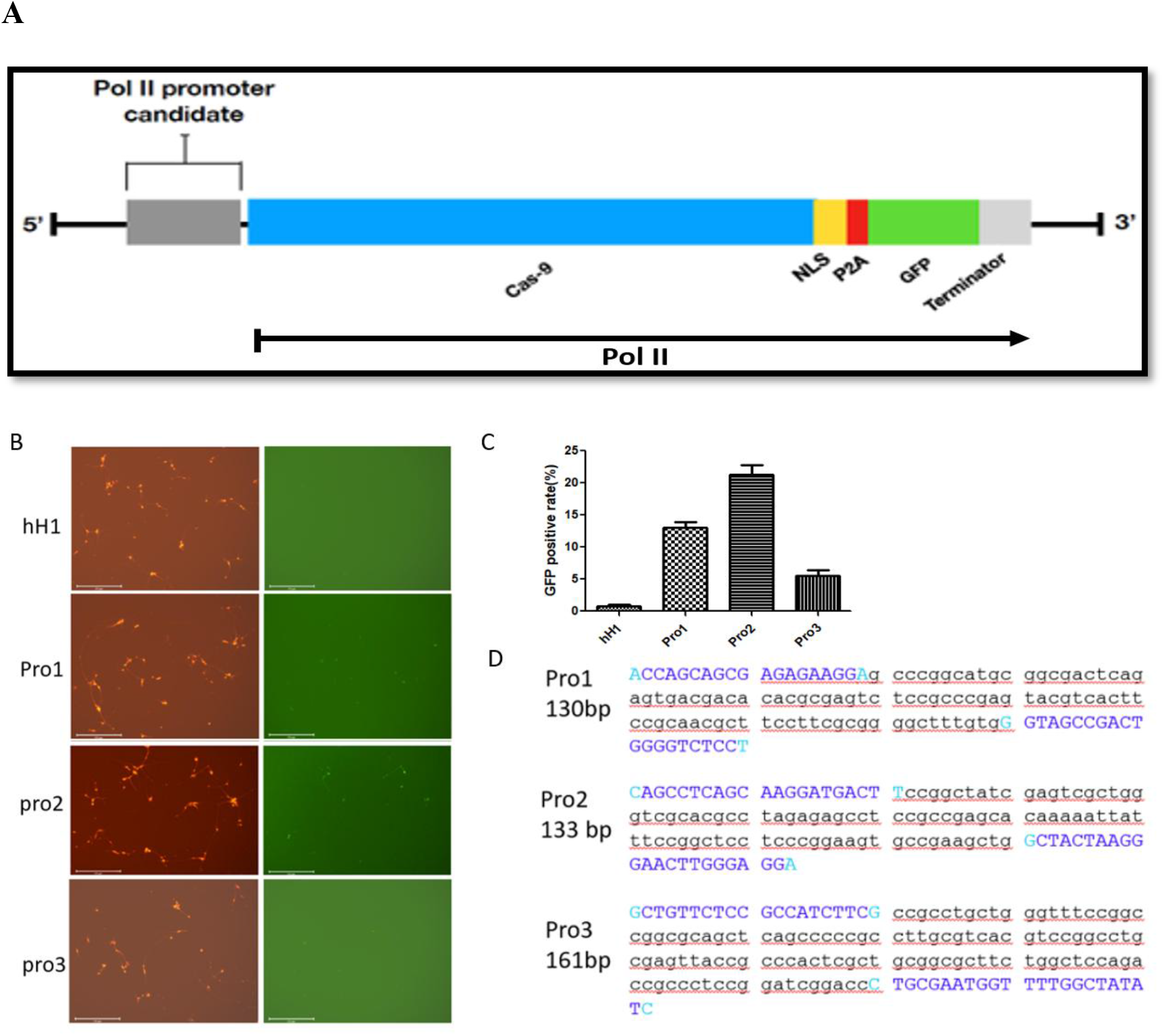
Screening for compact Pol II promoters in ESC-derived human RGCs using plasmid transfection. (A) Schematic representation of constructs designed to test candidate Pol II promoters for chimeric promoter generation. (B) Assessment of promoter activity in human RGCs. Left: tdTomato^+^ cells differentiated from H9 ES cells under the endogenous BRN3B promoter, confirming RGC identity. Right: GFP^+^ cells (48 h) following transfection with Pro1, Pro2, or Pro3 promoter–driven SpCas9-eGFP expression cassettes. (C) Quantification of GFP^+^ hRGCs after transfection with Pro1–Pro3 promoter constructs. (D) Nucleotide sequences of the three highest-expressing promoters. Uppercase letters denote primer sequences used for promoter cloning, while lowercase letters indicate the remaining promoter sequences.

From this screen, the top three Pol II candidate promoters were selected based on their bidirectional activity, promoter size (<165 bp), exogenous GFP expression, and endogenous expression levels in RGCs (based on unpublished RNA-seq data). These candidates, along with the human H1 Pol III promoter, were cloned into a GFP reporter construct and individually transfected into hESC-derived RGCs. GFP expression analysis revealed that Promoter 2 (133 bp) exhibited the highest activity among the tested constructs, representing the most efficient compact bidirectional Pol II promoter (Figure 4B, 4C). The sequences of the top three candidate Pol II promoters are shown in Figure. 4D.

### Evaluating Human 7SK, H1, and U6 Pol III Promoter Activity Using a CRISPR/Cas9 Reporter System

The human U6, 7SK, and H1 promoters are commonly used Pol III promoters, with 7SK and H1 also functioning as bidirectional promoters [14]. Traditionally, Pol III activity is assessed using techniques such as Northern blotting. However, we employed a reporter gene editing system in combination with our Pol II–Pol III chimeric constructs (sharing the same Pol II element) to evaluate Pol III promoter function (Figure 5A).

**Figure 5.**
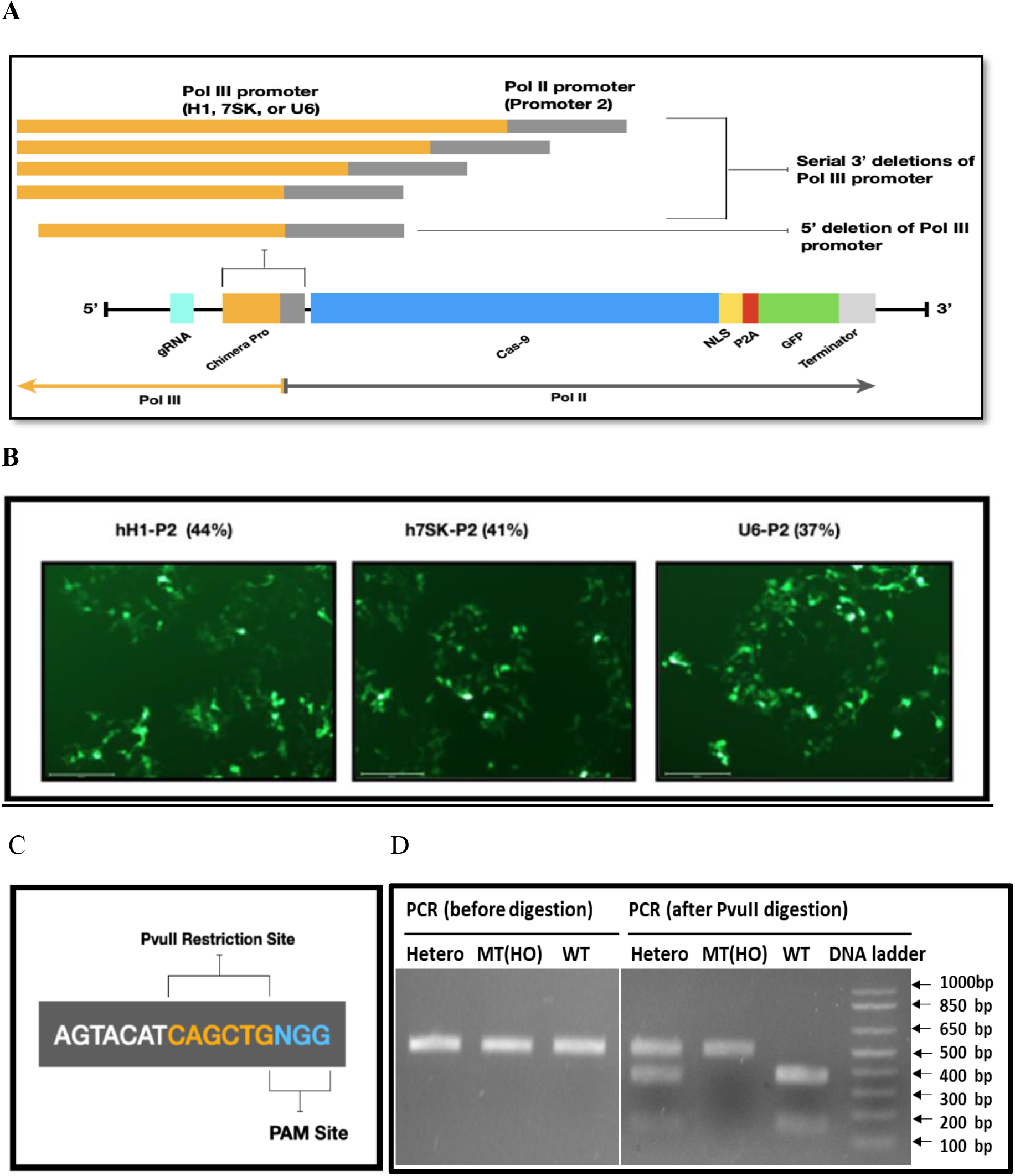
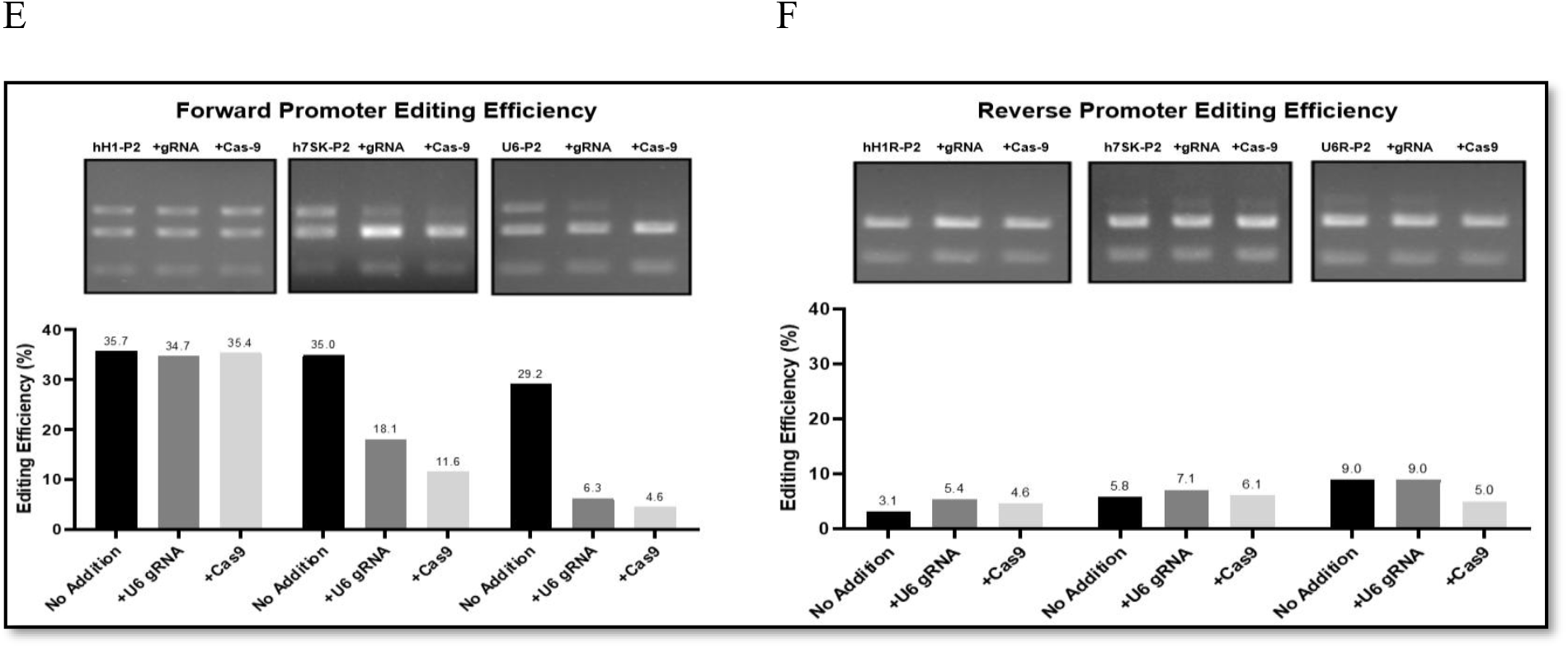
Hybrid promoter analysis reveals unidirectional Pol III activity of human U6, 7SK, and H1 promoters. (A) Schematic representation of Pol II–Pol III hybrid promoters and serial deletions used for functional analysis. (B) Fluorescence microscopy images show GFP-positive HEK 293 cells transfected with constructs containing the human H1 (hH1-P2), 7SK (h7SK-P2), or U6 (U6-P2) promoters in the P2 orientation. The percentage values in parentheses indicate the proportion of GFP-positive cells 48 hours after plasmids transfection. (C) Schematic of the target sequence highlighting the PvuII restriction site (orange) adjacent to the protospacer-adjacent motif (PAM, blue). (D) Validation of CRISPR/Cas9-mediated editing using a PvuII restriction enzyme mapping assay. Agarose gel electrophoresis showing PCR amplicons from MT (homozygous mutant), WT (wild type), and Hetero (heterozygous mutant) samples before and after PvuII digestion. Loss or retention of the PvuII site indicates successful genome editing. The expected bands are 509 bp for MT (cut site was destroyed), 358 and 151 bp for WT (intact PvuII cut site), and 509, 358 and 151 bp for Hetero. (E, F) Representative gel electrophoresis results showing CRISPR/Cas9-mediated editing with human H1, 7SK, and U6 promoters in the forward (E) and reverse (F) orientations. Editing reactions were performed with the order as follows: with no additional plasmid transfection, with additional gRNA plasmid transfection and with additional SpCas9 plasmids transfection, Quantification of editing efficiency (%) for each promoter and orientation based on band intensities from the gels above.

Using this system, we first tested the activity of the most widely used Pol III promoters. The reporter includes a PvuII restriction site located immediately upstream of the SpCas9 NGG PAM site. In the unedited (wild-type) sample, PvuII-HF cleaves the DNA at its recognition site, producing two fragments of ∼358 bp and ∼151 bp. If gene editing occurs, the Cas9-gRNA complex cleaves the genomic DNA and disrupts the PvuII site, preventing digestion. In this case, gel electrophoresis reveals a single intact 509 bp fragment.

In many cases, three bands were observed: the uncut 509 bp fragment and the two smaller digested fragments (358 and 151 bp), indicating a mixture of edited and unedited DNA. Editing efficiency was determined by quantifying the relative band intensities. It’s important to note that the measured editing efficiency may be limited by the transfection efficiency, even when editing reaches its maximum potential. In our system, gene editing efficiency was approximately 36%, which corresponded closely with the observed transfection efficiency in HEK293A cells (Figure 5B, 5C). Promoter 2 also reached maximum efficiency of Pol II function because additional Cas9 plasmids did not improve the reporter gene editing efficiency (Figure 5E). All three of these promoters appeared to have similar maximal Pol III promoter strength, since additional U6-gRNA plasmids did not improve the reporter gene editing efficiency (Figure 5E). We then tested whether these Pol III promoters have bidirectional Pol III function by inverted insertions and fused with the Pol II promoter but found that all these promoters lost Pol III function with inversion (Figure 5F).

### Optimization of Minimal Pol III Promoter Size through Serial Deletions and Functional Assessment of Hybrid Promoters

The original lengths of the U6, 7SK, and H1 Pol III promoters are 257 bp, 241 bp, and 233 bp, respectively (Figure S1). When fused with Promoter 2 (133 bp), a compact Pol II element, the resulting chimeric Pol II–Pol III promoters were approximately 400 bp in length—still larger than the desired size for efficient delivery vectors.

To reduce the overall length, we performed serial deletions from the 3′ end of each Pol III promoter sequence. These truncated versions were fused to Promoter 2 to generate compact hybrid promoters. Deletions were made upstream of any known proximal sequence elements and TATA boxes to preserve essential Pol III functionality.

Our results showed that deletion of 76 bp and 90 bp from the 3′ end of the H1 promoter retained most of its Pol III activity. However, a larger deletion of 110 bp from the H1 promoter significantly reduced Pol III function by approximately 80% with combined 5′ and 3′ truncations nearly abolishing pol II function. This indicated that the minimal hH1 promoter length is 143bp with retaining Pol III function (Figure 6A). Similarly, Stepwise 3′ deletions (d59, d80) lowered activity, and combined 5′ and 3′ deletions nearly eliminated function of the 7SK (Figure 6B). The minimal h7SK promoter length is 161 bp with retaining most its Pol III function. Truncations at the 3′ end (d52, d67) retained U6 Pol III activity, with combined 5′ and 3′ deletions almost completely abolishing activity. The minimal hU6 promoter length is 190 bp (Figure 6C). Interestingly, while stepwise deletions of up to 90 bp from the 3′ end of the Pol III component retained Pol III activity, even small deletions (9–12 bp) from the 5′ end of the Pol III sequence abolished its function (Figure 6A–C).

**Figure 6.**
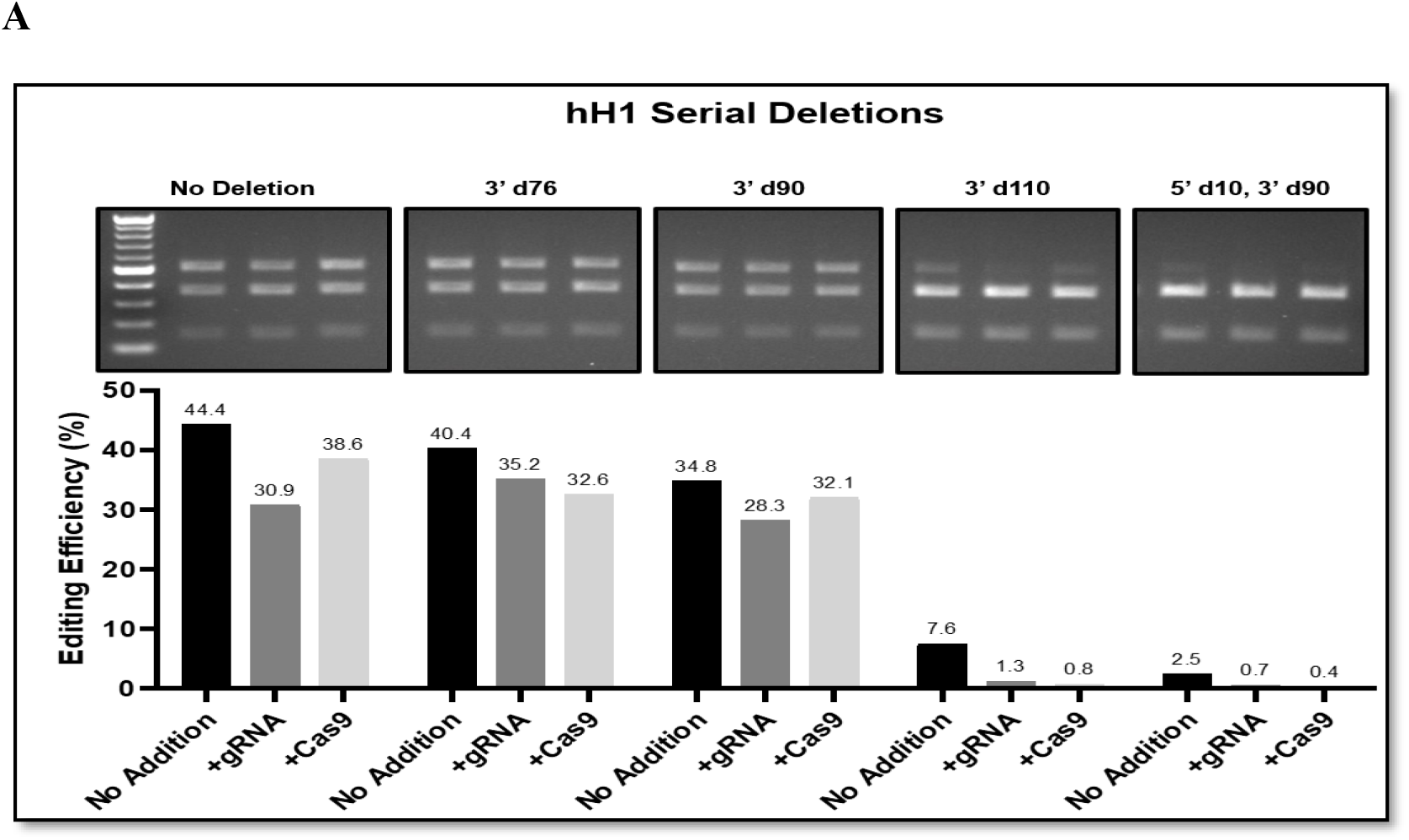
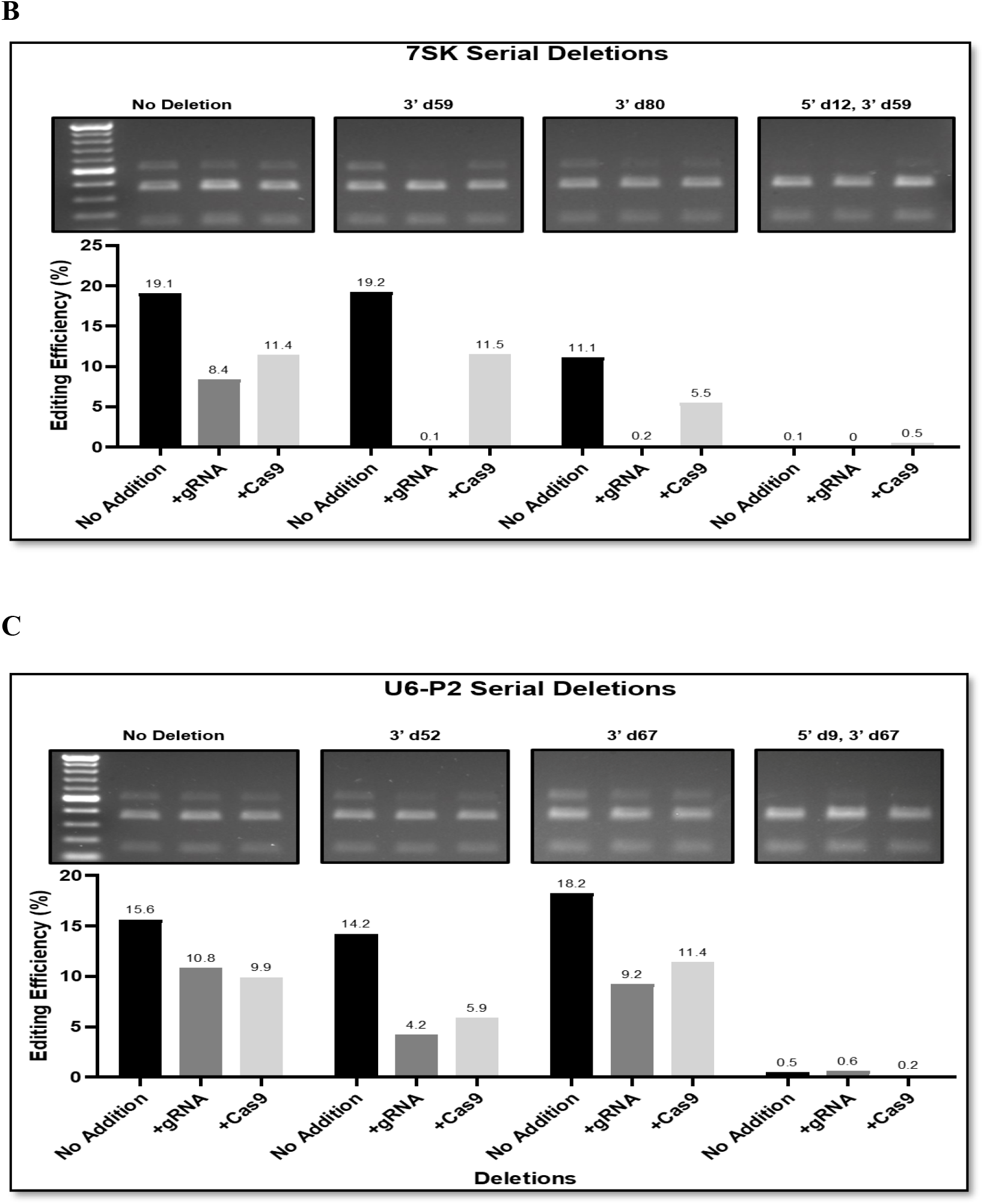
Determination of minimal Pol III promoter lengths in hybrid constructs using a CRISPR/Cas9 reporter assay. HEK293 cells were transfected with hybrid constructs, and genomic DNA was extracted 48 hours post-transfection for gene editing analysis (as described in Figure 5). Hybrid Pol II–Pol III constructs containing the human H1 (hH1), 7SK (h7SK), or U6-P2 (hU6-P2) promoters were subjected to serial deletions to identify the minimal sequences required for Pol III transcriptional activity. “3′d” indicates deletions from the 3′ end of the promoter, and “5′d” indicates deletions from the 5′ region. Representative gel electrophoresis images (top) show CRISPR/Cas9 reporter assay products, and corresponding bar graphs (bottom) display editing efficiency (%) under no-addition, +gRNA, or +Cas9 conditions. (A) Minimal length determination of the hH1 promoter. (B) Minimal length determination of the h7SK promoter. (C) Minimal length determination of the hU6 promoter.

These findings highlight the critical sequence requirements at the 5′ end of Pol III promoters and demonstrate a viable strategy for optimizing the size of dual Pol II–Pol III hybrid promoters. Thus, the chimera promoters hH1del90-Pro2, h7SK-del80-pro2, and U6-del67-Pro2 (276, 294, and 323 bp, respectively) were functional like their original chimera promoters (hH1-pro2, h7SK-pro2 and hU6-pro2).

### Cell-Type Preferential Expression and Gene Editing in Purified Human RGCs using Optimized Hybrid Promoters

We next assessed whether the optimized hybrid promoters could drive effective Cas9 and gRNA expression in both human HEK293 cells and hESC-derived retinal ganglion cells (RGCs). As an initial step, we determined the maximum gene editing efficiency achievable in HEK293 cells. Reporter gene editing at the DLK locus reached ∼35–40%, closely matching the plasmid transfection efficiency in these cells. Since gene editing efficiency can be constrained by vector delivery, fluorescence-activated cell sorting (FACS) was used to enrich for GFP-positive (successfully transfected) cells.

After fluorescence-activated cell sorting (FACS) enrichment of GFP-positive HEK293A cells, reporter gene editing efficiencies reached almost 100.00% for constructs driven by the hH1-del90-Pro2 and U6-del67-Pro2 hybrid promoters, indicating near-complete editing among transgene-expressing cells. In contrast, the h7SK-del80-Pro2 construct achieved a substantially lower but measurable editing efficiency of 55.1%, suggesting promoter-dependent differences in CRISPR activity. Importantly, no detectable editing (0%) was observed in the GFP-negative cell populations across all three constructs, confirming that genome editing was strictly dependent on transgene expression rather than nonspecific background activity (Figure 7A, 7C).

**Figure 7.**
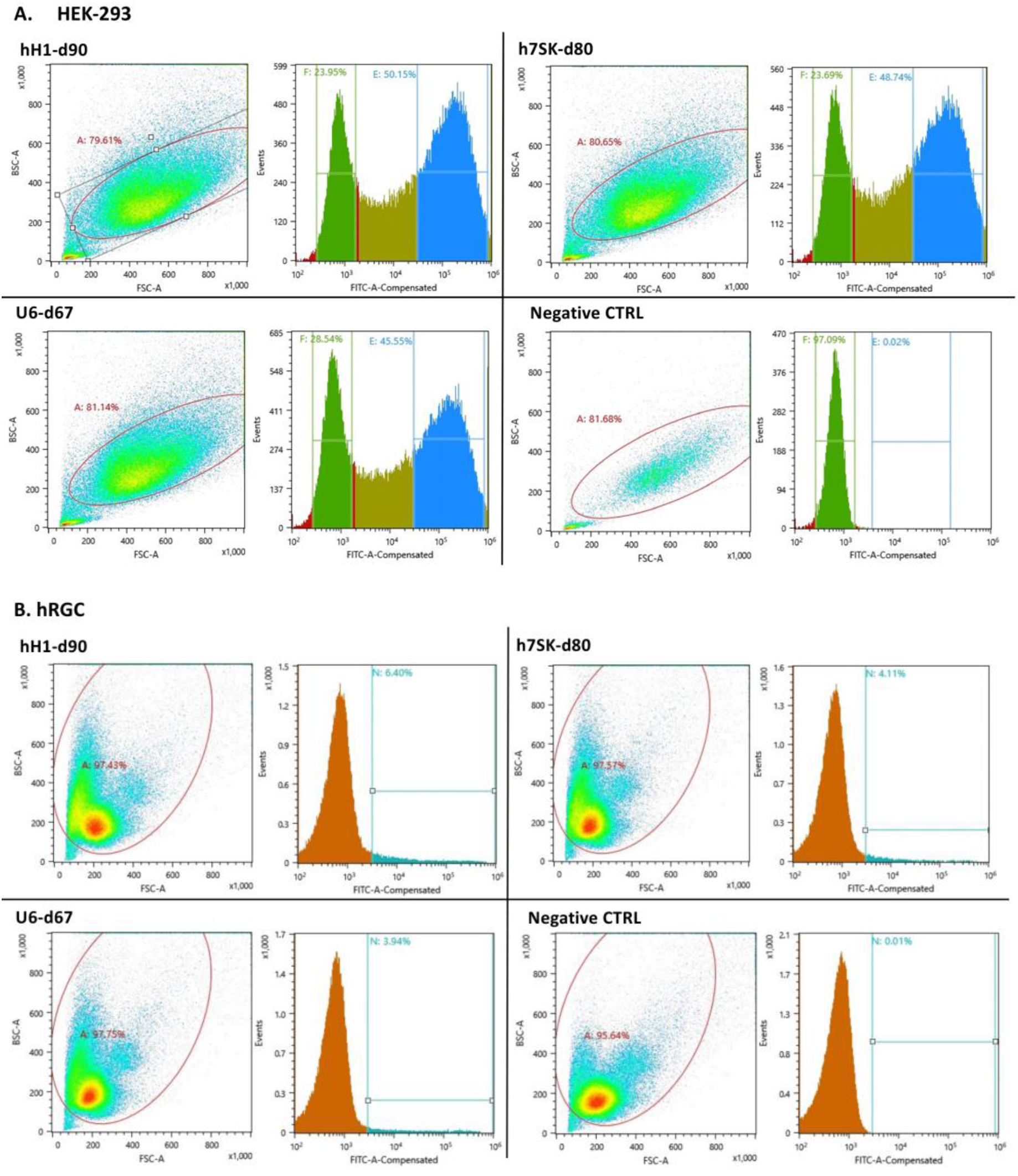
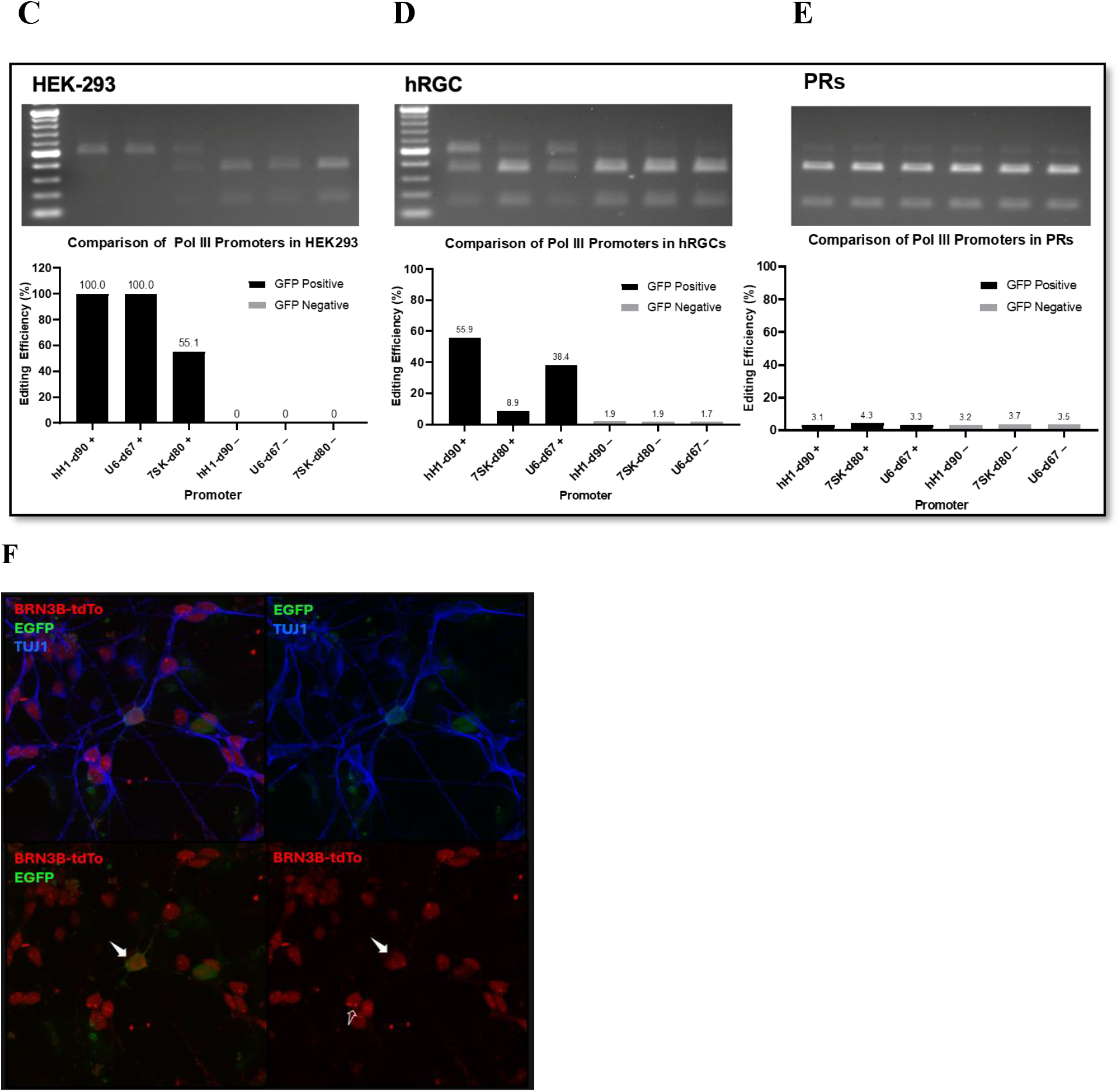
CRISPR/Cas9 editing in FACS-purified GFP^+^ HEK293A/hRGCs and validation of reporter overlap with endogenous BRN3B expression. (A, B) GFP^+^ HEK293A and hRGCs transfected with hH1-del90-Pro2, h7SK-del80-Pro2, or U6-del67-Pro2 plasmids were FACS-purified 48 h after transfection. Only highly GFP^+^ cells were used for reporter editing analysis. (C–E) Reporter gene editing efficiencies in purified HEK293A, hESC-derived RGCs, and hESC-derived photoreceptors from retinal organoids were quantified from band intensities in the gels shown above. (F) hESC-derived retinal organoid RGCs (day 80) were analyzed for co-expression of endogenous BRN3B-driven tdTomato and exogenous EGFP. (Top left) TUJ1 (blue) marks neurons, tdTomato (red) reports BRN3B activity, and EGFP (green) indicates exogenous reporter expression. (Top right) TUJ1 (blue) and EGFP (green) mark differentiated RGCs and exogenous expression activity. (Bottom left) Co-localization of tdTomato and EGFP in RGCs. (Bottom right) tdTomato expression under the BRN3B promoter. (white boxes) highlight cells co-expressing tdTomato and EGFP (arrows), confirming reporter activity in BRN3B^+^ RGCs. Scale bars, 100 µm.

We next evaluated the performance of these hybrid promoter constructs in sorted GFP-positive hESC-derived retinal ganglion cells (RGCs) to assess their functionality in a disease-relevant neuronal context. Editing efficiencies in GFP-positive RGCs reached 55.9% for hH1-del90-Pro2, 8.9% for h7SK-del80-Pro2, and 38.4% for U6-del67-Pro2, demonstrating robust but promoter-specific editing activity in differentiated human neurons. In contrast, GFP-negative RGC populations exhibited markedly reduced editing efficiencies of 1.9%, 1.9%, and 1.7%, respectively, indicating strong enrichment of editing activity within transgene-expressing cells. These differences highlight both the effectiveness of GFP-based enrichment and the relative specificity of the hybrid promoters in driving genome editing in hESC-derived RGCs (Figure 7B, 7D, 7F).

To evaluate promoter activity in photoreceptor cells, we used H9 knock-in reporter ESCs expressing tdTomato under control of the cone-rod homeobox (CRX) promoter (H9-CRX), which were differentiated into 3D retinal organoids. After dissociation and reseeding, the cells were transfected with either hH1-del90-Pro2::GFP or CMV-GFP plasmids. In the CMV-GFP group, GFP expression was observed in tdTomato-positive photoreceptor cells, indicating strong promoter activity in this lineage. However, fewer tdTomato-positive cells expressed GFP following transfection with the hH1-del90-Pro2 plasmid, suggesting that this hybrid promoter has very limited activity in photoreceptors (Figure 7D).

## Discussion

Gene therapy is a molecular approach that treats diseases by correcting defective genes or introducing therapeutic ones. A critical step in this process involves incorporating gene-editing components or therapeutic genes into a delivery vector, along with a promoter to control expression. The promoter plays a key role in determining both the cell specificity and the expression level of the therapeutic protein encoded by the transgene. The CRISPR/Cas9 system is a powerful genome-editing tool and holds great promise for treating genetic diseases, especially gain-of-function disorders. In addition to therapeutic applications, gene-editing technologies are also invaluable for creating in vitro disease models, exploring disease mechanisms, and identifying therapeutic targets.

Earlier gene-editing platforms, such as zinc-finger nucleases (ZFNs) and transcription activator-like effector nucleases (TALENs), have been used for disease modeling and functional studies. However, CRISPR/Cas9 offers several advantages over ZFNs and TALENs, including simpler design, greater flexibility in target selection, and the ability to simultaneously modify multiple genomic loci [17–19]. Targeting gene expression to the appropriate cell types is crucial for the success of gene therapy. This involves not only optimizing delivery methods but also identifying suitable promoters that ensure cell type-specific expression. Viral vectors, such as adeno-associated virus (AAV), remain the most efficient delivery vehicles, but their broad tropism can lead to off-target expression in unintended cell types [17]. Using selective, cell-specific promoters—regulated by transcription factors and RNA polymerase activity—is an effective strategy to restrict transgene expression to desired cell populations. The use of AAV vectors imposes strict size limitations on gene therapy constructions, making the development of compact promoters essential. This challenge is particularly significant for neurological disorders, where precise delivery and expression of gene-editing tools in post-mitotic neurons, such as retinal ganglion cells (RGCs), is required and can be difficult to achieve [20, 21].

In this study, we introduced a novel compact promoter selection strategy based on our observation that exogenous transgene expression driven by compact promoters does not strongly correlate with endogenous gene expression. This finding suggests that comparing transgene expression levels among different cell lines is a more effective method for selecting compact promoters than relying on endogenous expression patterns. Using this strategy, we screened 509 bidirectional promoters and identified three top candidates, ultimately isolating a 133 bp Pol II promoter (Promoter 2) capable of driving exogenous expression in human RGCs.

Previous work by Nieuwenhuis et al. compared the activity of five different promoters in mouse RGCs using AAV delivery and identified the SYN promoter as a strong candidate [13]. However, its relatively large size (∼700 bp) makes it less suitable for single-AAV gene-editing applications, which require compact elements due to vector packaging constraints.

To overcome this limitation, we engineered Pol II–Pol III chimeric promoters by fusing our compact 130 bp Pol II promoter with truncated versions of three commonly used Pol III promoters (U6, H1, and 7SK). Through systematic 3′ end deletions and reporter gene assays, we generated three compact chimeric promoters that retain strong Pol III activity while meeting the size requirements for single-AAV packaging. These constructions offer new tools for single-vector gene therapy approaches targeting human RGCs.

The gene-editing capability of these hybrid promoters was validated in human RGCs derived from pluripotent stem cells. Furthermore, we demonstrated that these promoters exhibit preferential activity in RGCs compared to photoreceptors in retinal organoid cultures. However, their activity has not been evaluated in other retinal cell types, such as bipolar, amacrine, horizontal, or Müller glial cells. Additionally, in vivo validation of the single-AAV constructs was not performed, which represents a limitation of the current study.

