## supplementary file for "Identification a Compact Promoter using a New Promoter Selection Strategy and Engineering Hybrid Pol II/III Enable Efficient Genome Editing in Human Retinal Ganglion Cells"

### Supplemental material

1. Sequence of three commonly used pol III promoters and minimized promoters:

1)Human H1 original promoter (233bp)

GGTGGAATTCGAACGCTGACGTCATCAACCCGCTCCAAGGAATCGCGGGCCCAGTGTCACT
AGGCGGGAACACCCAGCGCGTGCGCCCTGGCAGGAAGATGGCTGTGAGGGACAGGGGAG
TGGCGCCCTGCAATATTTGCATGTCGCTATGTGTTCTGGGAAATCACCATAAACGTGAAAT
GTCTTTGGATTTGGGAATCTTATAAGTTCTGTATGAGACCAC**T**CTTTCCCT

2)hH1 minimized promoter with retaining function (143bp, del90bp)

TGGCAGGAAGATGGCTGTGAGGGACAGGGGAGTGGCGCCCTGCAATATTTGCATGTCGCTA
TGTGTTCTGGGAAATCACCATAAACGTGAAATGTCTTTGGATTTGGGAATCTTATAAGTTC
TGTATGAGACCAC**T**CTTTCCC**T**

2. Human 7SK promoter

1)Original size is 241bp

CTGCAGTATTTAGCATGCCCCACCCATCTGCAAGGCATTCTGGATAGTGTCAAAACAGCCGGAAAT
CAAGTCCGTTTATCTCAAACTTTAGCATTTTGGGAATAAATGATATTTGCTATGCTGGTTAAATTA
GATTTTAGTTAAATTTCCTGCTGAAGCTCTAGTACGATAAGTAACTTGACCTAAGTGTAAAGTTGA
GATTTCCTTCAGGTTTATATAGCTTGTGCGCCGCCTGGGTACCTC

2)H7SK minimized promoter with retaining function (161bp, del80 bp)

AAACTTTAGCATTTTGGGAATAAATGATATTTGCTATGCTGGTTAAATTAGATTTTAGT
TAAATTTCCTGCTGAAGCTCTAGTACGATAAGTAACTTGACCTAAGTGTAAAGTTGAGA
TTTCCTTCAGGTTTATATAGCTTGTGCGCCGCCTGGGTACCTC

3. U6 promoter

1)Original size is 257bp

AAGGTCGGGCAGGAAGAGGGCCTATTTCCCATGATTCCTTCATATTTGCATATACGATACAAGGC
TGTTAGAGAGATAATTAGAATTAATTTGACTGTAAACACAAAGATATTAGTACAAAATACGTGAC
GTAGAAAGTAATAATTTCTTGGGTAGTTTGCAGTTTTAAAATTATGTTTTAAAATGGACTATCAT
ATGCTTACCGTAACTTGAAAGTATTTCGATTTCTTGGCTTTATATATCTTGTGGAAAGGACG

2)hU6 minimized promoter with retaining function (190bp, del67bp)

TTAGAGAGATAATTAGAATTAATTTGACTGTAAACACAAAGATATTAGTACAAAATACGTGACGT
AGAAAGTAATAATTTCTTGGGTAGTTTGCAGTTTTAAAATTATGTTTTAAAATGGACTATCATAT
GCTTACCGTAACTTGAAAGTATTTCGATTTCTTGGCTTTATATATCTTGTGGAAAGGACG
